# Predicting gene regulatory network interactions with high certainty using the Linear Profile Likelihood - LiPLike

**DOI:** 10.1101/2025.03.24.644951

**Authors:** Rasmus Magnusson

## Abstract

Gene regulatory network inference typically revolves around the use of expression data to predict the most likely underlying network, and these predictions of transcription factor-target regulations have a low accuracy and are full of false positives. This low accuracy is in part due to a high degree of correlation between regulatory transcription factors. However, most gene regulatory network inference methods aim at predicting the maximizing a likelihood function, overlooking the degrees of freedom in almost equally likely alternative solutions. This chapter introduces the LInear Profile LIKE-lihood, LiPLike, which controls for this uncertainty by only selecting network interactions that are uniquely needed to explain the data. If there exists an alternative solution B to an interaction prediction A, LiPLike predicts neither A nor B, whereas state-of-the-art inference methods typically predict either A or B, or both. LiPLike can be used as a standalone inference method, or to stratify the predictions of alternative inference methods into sets of low and high confidence.

## 1 Background

Gene regulatory networks that are inferred from gene expression data will contain numerous false positive identifications, with blind tests in real biological data showing an accuracy substantially below 0.5 even in the top predictions of high-performing inference methods [15, 17]. One of the most pronounced culprits for this low performance is the occurrence of correlated explanatory variables, typically the gene expression of transcription factors [2, 6, 26]. Although most gene regulatory network inference (GRNI) methods aim to extract the most likely regulators of a downstream gene [28], most approaches pay little or no attention to the existence of alternative solutions, i.e., different combinations of regulators that explain the same downstream gene expression [8, 22, 26, 27, 30]. This chapter argues that an optimal GRNI method would also stratify predictions into layers of confidence, where the most certain predictions are uniquely needed to fit the data, as opposed to ordering predictions by maximizing the log likelihood.

This chapter will also discuss how analyses of parametric uncertainty are an integral part of techniques beyond GRNI, with the example shown being mechanistic modeling using ordinary differential equations (ODEs). It will briefly present the method of profile likelihood analysis and how it can be used to establish intervals of possible parameter values during inference.

Lastly, this chapter will present the LInear Profile LIKElihood (LiPLike), a GRNI tool built to focus on predicting interactions with high probability, as opposed to recreating full gene regulatory networks. The LiPLike approach is based on the observation that linear combinations of transcription factors often capture the main modes of regulation in gene expression data [28]. Therefore, a gene regulation link should be excluded from the network if an alternative linear combination of regulators can adequately explain the expression of a downstream gene. This ensures that only the most probable interactions are predicted (Fig. 1).

**Figure 1:**
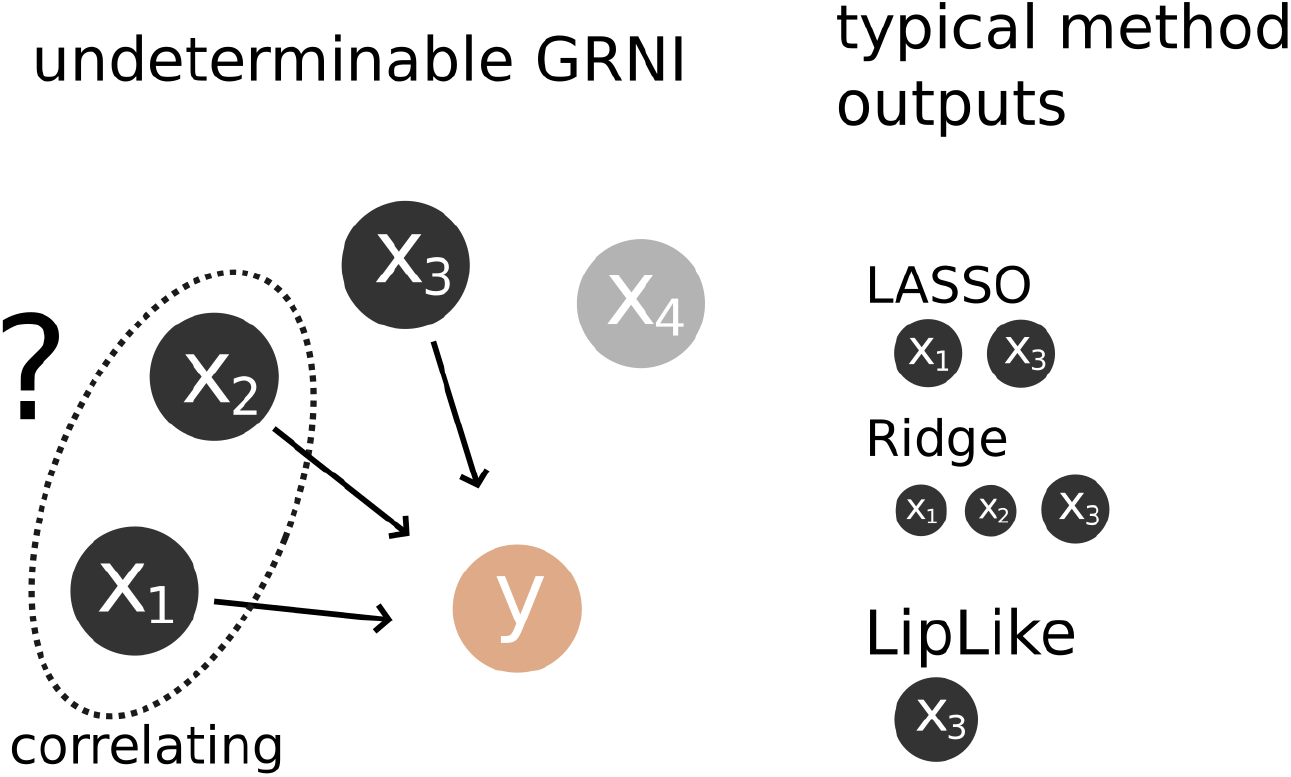
LiPLike rationale. Where there are two correlating explanatory variables, the corresponding GRNI problem is indeterminable. Depending on chosen model, predicted regulators of dependent variable *y* will typically include *X*_3_, and either *X*_1_ and *X*_2_, whichever minimizes the objective function.

### Rationale behind estimating parameter uncertainty in GRNI

Collinearity, i.e. highly correlated explanatory variables, is one of the most complicating factors in feature selection [3, 7, 19]. In the field of gene regulatory network inference, correlated explanatory variables will have widely different impacts depending on which inference method is used. Consider a hypothetical scenario where *m* samples of gene expression measurements from *n* transcription factors (TFs) 1..*n* and one dependent gene have been collected, such that *m < n*. In this case, TF_1_ and TF_2_ correlate to a large extent, and TF_3_ has slightly less explanatory power than TF_1_ and TF_2_ individually. Consider the commonly used penalized linear regression models: LASSO (Least Absolute Shrinkage and Selection Operator) [25] and Ridge regression. By studying the amplitude of the inferred coefficients, the two models will predict different gene regulation structures. The LASSO model will select TF_3_ together with either TF_1_ or TF_2_, depending on which coincidentally had the highest explanatory power [15, 30]. Ridge regression would assign approximately equal coefficients to TF_1_ and TF_2_ due to their high correlation, effectively sharing the explanatory weight between them [25]. In contrast, TF_3_, being less correlated with the others, provides unique information, resulting in a comparatively higher coefficient despite its slightly lower individual explanatory power. In other words, both of these approaches to identify regulators for downstream target genes would fail to adequately capture the inherent uncertainty of whether TF_1_, TF_2_, or both are potential regulators.

This uncertainty is inherent in most GRNI methods. Yet, extracting high-confidence predictions from broader regulatory estimates is often overlooked. Indeed, the majority of network inference methods will maximize a likelihood function relating to network inference, often at the expense of a large number of false positive predictions. This aspect is perhaps best illustrated by the GRNI challenge Dialogue on Reverse Engineering Assessment and Methods 5 (DREAM5) [17], where participants aimed to predict networks for four datasets with hidden network structures from independent sources. In DREAM5, the focus was to recreate the most likely underlying network from data, yet, the resulting publications did not discuss the correlaton of gene expression among transcription factors. [15, 17]

### Estimating parameter uncertainty in mechanistic models

LiPLike draws inspiration from parameter identifiability analyses in mechanistic modeling [5, 9, 12, 16]. In this field, analyses typically focus on a limited number of variables with known or hypothesized interactions. Furthermore, there is most often some prior knowledge of how these variables interact, which is formulated into a set of hypotheses [16, 21]. Next, models can be rejected by their ability to explain data, which is typically tested using heuristic search algorithms of parametric space, necessitated by non-linear dependencies and non-convex likelihood surfaces [4]. In Figure 2, and the corresponding equations 1-3, a small toy mechanistic model is shown.

**Figure 2:**
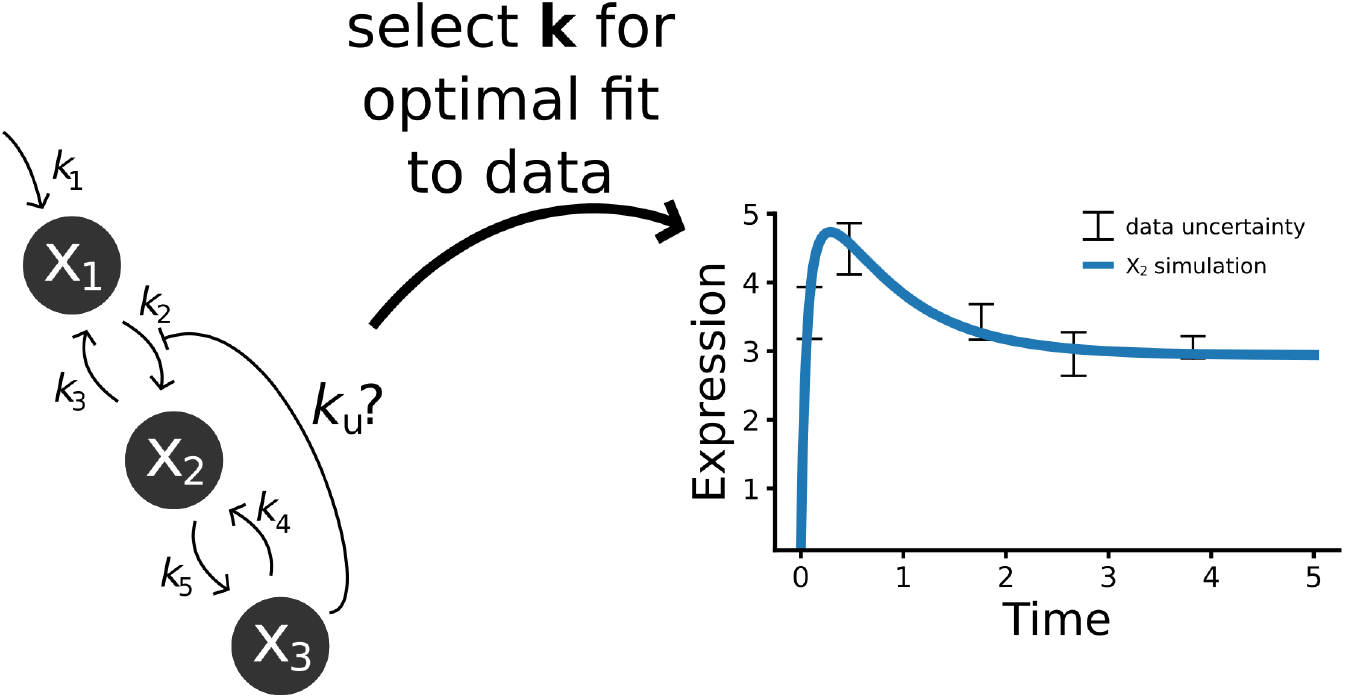
Simple toy mechanistic model with unknown parameters being optimized to fit to data. A model hypothesis is formulated, which typically includes several unknown parameters, here shown as (*k*_1_..*k*_5_, *k*_*u*_). These parameters are estimated by optimizing the likelihood function, often requiring heuristic search algorithms and numerical integration for non-convex problems.

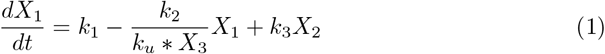

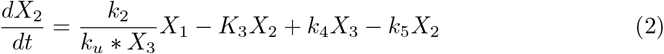

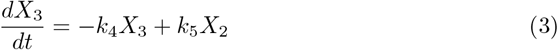

In 1-3, *X*_1_-*X*_3_ denote the states of the model, while *k*_1_ −*k*_5_, *k*_*u*_ represent the unknown parameters. The model describes the reaction rates of how *X*_1_ is broken down into *X*_3_, and the model can be numerically integrated to compare with the data.

Arguably, the scientific value of this approach lies in the ability to reject model structures, which can be done in several ways. As such, a model can be rejected by the inability to fit existing observations, where these observations are either qualitative, for example, *Can the model reproduce the observed overshoot in the data?*, or the rejection can be quantitative, e.g. *Are the residuals between model output and observed data too large to result from white noise?*.

From the perspective of parametric uncertainty in mechanistic modeling, the most relevant analysis is arguably the profile likelihood [9, 23]. The profile likelihood assesses how the model’s fit varies with changes in a specific parameter. In Figure 3, a figurative behavior of the model fit with respect to given values of *k*_*u*_ is shown. To derive the likelihood profile of parameter *k*_*u*_, the remaining model parameters are iteratively reestimated given a range of set values of *k*_*u*_, and the best model fits to data are recorded, here as the residual sum of squares (RSS) weighted by the uncertainty of data. The profile likelihood of a parameter *k*_*u*_ can be expressed as an analysis of the negative logarithmic likelihood (−*LL*) of the model residuals with respect to different values of ζ, as expressed in (4)

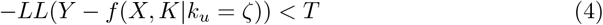

**Figure 3:**
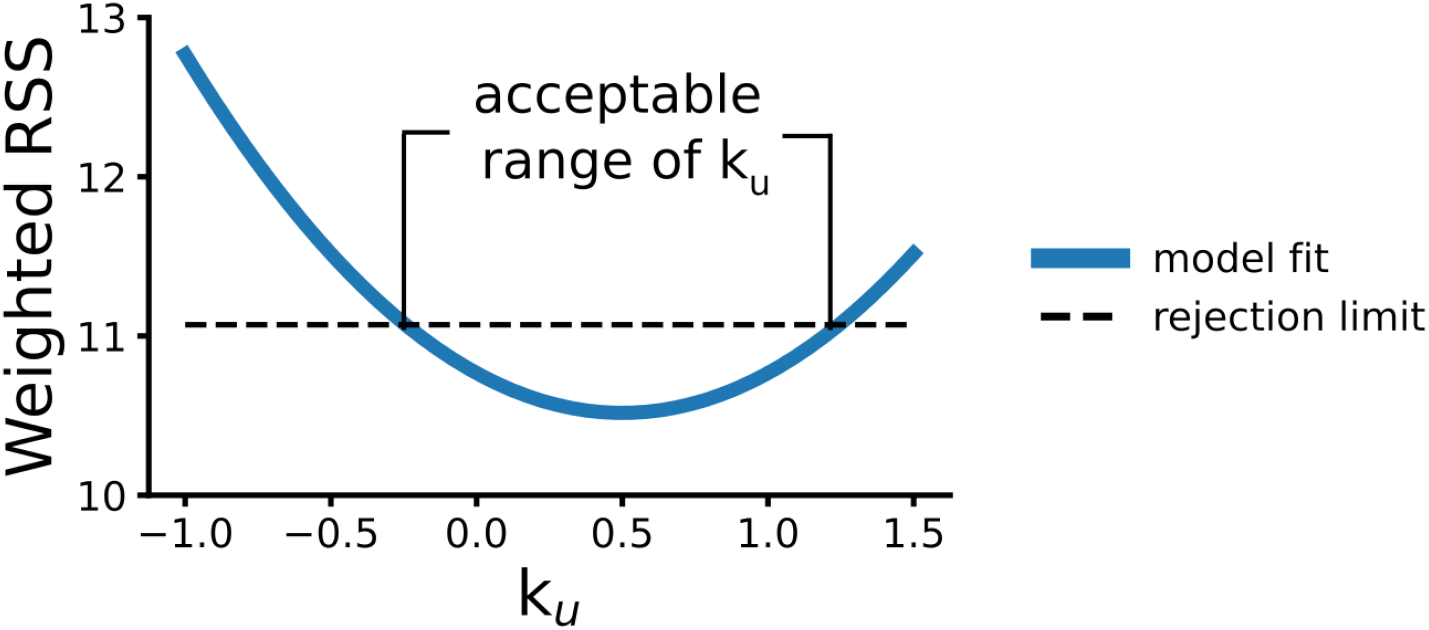
Estimating parametric uncertainty of *k*_*u*_ in a profile likelihood. By analyzing the model’s optimal fit to data with respect to a given value for *k*_*u*_, while re-estimating the rest of the parameters, a range of acceptable values for *k*_*u*_ can be determined. Shown is a schematic of a profile likelihood, showing that −0.25 *< k*_*u*_ *<* 1.2.

In (4), the fit between the observed data *Y* and the model prediction *f* as a function of the states *X* and parameters *K* is calculated. Applied to the parameter *k*_*u*_, different values ζ are assigned to *k*_*u*_, and the optimal model fit is calculated. Acceptable values of *k*_*i*_ are those for which the −*LL*, evaluated at ζ, falls below a threshold *T*, typically derived from a *χ*^2^-test [16].

## 2 Materials

LiPLike is a computational method for inferring gene regulation interactions with high certainty. As we will see in the methods section, the computational implementation of LiPLike is surprisingly straightforward and can be implemented with a few lines of code in most programming languages. Nevertheless, LiPLike is available as a Python implementation, with only Numpy and Scikit-learn as dependencies.

The implementation of LiPLike for gene regulation inference can be summarized as follows;

1. Extracting **gene expression** data. LiPLike assumes a linear model, and it is recommended to perform appropriate normalizations.
2. Separating the data into regulators and target genes. Typically, the data are divided into a set of transcription factors and target genes, which form the independent and dependent variables, respectively. **Thus, a defined set of TFs is needed**.
3. Limiting the explanatory variables (optional). As the rationale of LiPLike is to not predict any interaction where there is an alternative linear combination of regulatory elements that can explain the expression of the target gene, highly correlated independent variables will likely not be identified by LiPLike. **To adjust for this, it is possible to merge TF isoforms into their respective mean values**.
4. LiPLike prioritizes accuracy over recall. Therefore, this method is not well-suited for predicting the full underlying network. **A prominent aspect is that LiPLike can instead be used to stratify the results of other GRNI algorithms, such as the LASSO [25], the ElasticNet [30], ARACNE [18], TIGRESS [7], or the Inferelator [13]**.

## 3 Methods

### Using the LiPLike to extract gene regulatory elements of high confidence

In GRNI methods, an important question is whether the parameters of the model can be reliably identified. Whereas the profile likelihood operates under a fixed model structure, GRNI methods are typically implemented with considerable flexibility in model formulation.

The LiPLike GRNI tool was developed to explicitly account for the existence of alternative regulatory solutions [15]. Drawing inspiration from the profile likelihood (as shown in Figure 3), LiPLike compares the fit of gene expression for a target gene *y* using the full set of explanatory variables *X* to that obtained when excluding a single variable *X*_*i*_. If removing *X*_*i*_ significantly worsens model fit, the interaction was uniquely needed to explain the gene expression of target gene *y*. This approach is demonstrated in Eq. 5-6.

While the rationale behind LiPLike, along with several aspects of its implementation, may be relevant to other approaches, the core methodology of LiPLike is built upon a linear model describing the relationship between regulators and target genes. This commonly assumed relationship is formalized in Equation 5:

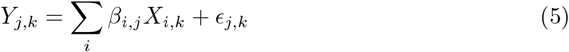

where *Y*_*j,k*_ is the expression of the *j*-th target gene in the sample *k, X*_*i,k*_ is the expression of the *i*-th regulator, and *β*_*i,j*_ ∈ R quantifies the contribution of *X*_*i*_ to *Y*_*j*_. The residual *ϵ*_*j,k*_ captures unexplained variation.

Next, the two relevant points of interest are where a) the full model is used to minimize the RSS, and b) where the interaction studied between the *i*-th regulator and the *j*-th dependent gene takes the value of 0 (Fig. 4). The importance of the interaction between the *i*-th regulator and *j*-th target gene is characterized by:

**Figure 4:**
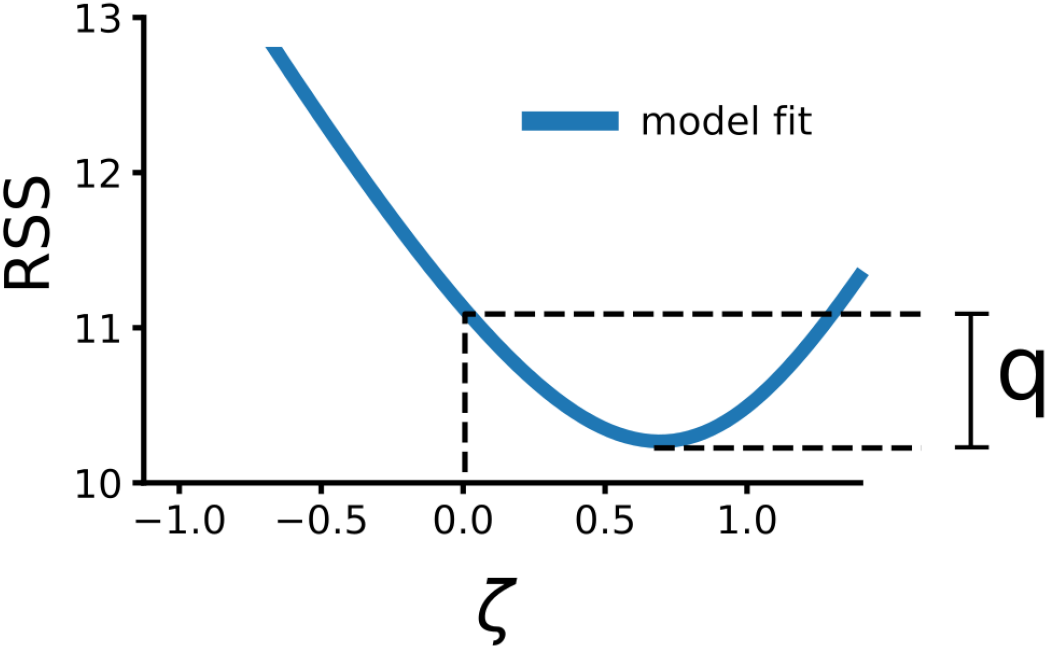
Schematic of extracting the LiPLike statistic *q*. The residual sum of squares (RSS) is shown as a function ζ, highlighting points where the full model minimizes the RSS and where the interaction is constrained to zero.

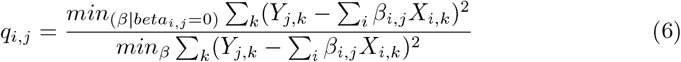

Here, *q*_*i,j*_ quantifies how much the RSS increases when the corresponding interaction is removed. Higher *q*_*i,j*_ values indicate that the interaction cannot be compensated for by other regulators. We note that *q >* 1. This approach identifies uniquely determined interactions, facilitating the construction of a high-confidence core of predictions in GRNI.

### Penalized LipLike models for undetermined problems

An attentive reader may notice that the linear model in Eqs. 5–6, which can be solved using ordinary least squares, assumes an overdetermined system. Specifically, when the number of samples *k* exceeds the number of predictors i (assuming a full-rank matrix *X*), the system is overdetermined, resulting in residuals greater than zero. In this scenario, LiPLike can be implemented directly.

However, when *i* ≤ *k*, the system becomes either exactly determined (*i* = *k*) or underdetermined (*i < k*). In these cases, the model often exhibits a perfect fit, causing the denominator in Eq. 6 to evaluate to zero, leaving the LiPLike statistic *q* undefined. To overcome this issue, the LiPLike software [15] incorporates an optional *L*_2_ regularization term. This regularization adds a penalty proportional to the sum of the squared coefficients *β* controlled by a penalty parameter 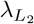. This term discourages large coefficient values, stabilizing the solution in underdetermined settings. Selection of 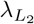 is recommended via cross-validation, using the full network to assess the denominator in Eq. 6. This approach ensures a balanced trade-off between model fit and regularization. In detail, the LiPLike algorithm now solves Equation 7:

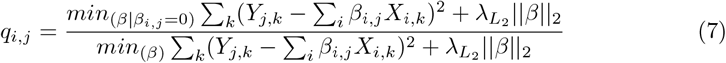

Here, 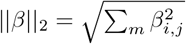 is the *L*_2_ norm of the coefficients, and 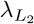 controls the strength of the penalty. By penalizing large coefficients, the solution avoids overfitting in underdetermined systems. Overall, this regularization ensures that the LiPLike statistic remains defined and stable, even when the system lacks sufficient constraints.

### Applying LiPLike in GRNI

The primary advantage of LiPLike is its ability to identify a core set of high-confidence gene-gene interactions. It is particularly effective as a complement to existing GRNI tools, enhancing the reliability of their predictions. In Magnusson and Gustafsson (2020) [15], LiPLike was applied to the DREAM5 GRNI challenge datasets [17]. The results demonstrated a significant improvement in prediction accuracy compared to all participating methods, including the community consensus model introduced by Marbach et al. [17]. In other words, compared to the top predictions from other methods in the DREAM5 challenge, LiPLike produced fewer false positive gene-gene interaction predictions [15].

It is important to note that LiPLike is intentionally designed to prioritize accuracy over recall. This focus ensures that the identified interactions are highly reliable, even if some true interactions are left unidentified. Therefore, LiPLike is best used to stratify predictions from independent GRNI methods into high- and lower-confidence sets (Fig. 5), providing a clearer framework for downstream biological interpretation. Magnusson and Gustafsson [15] employed this strategy and found that the intersection between community-based predictions and LiPLike results yielded substantially higher accuracy compared to either approach alone.

**Figure 5:**
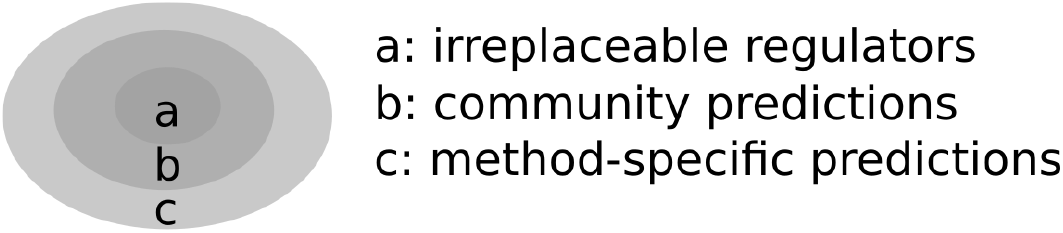
Using LiPLike for Prediction Stratification. While LiPLike prioritizes accuracy, it does so at the expense of recall. To balance this trade-off, it is recommended to use LiPLike in combination with other GRNI methods. This stratified approach classifies predictions based on confidence levels: method-specific predictions have lower confidence, while those supported by multiple methods are more reliable. Irreplaceable interactions, uniquely identified by LiPLike and corroborated by other methods, form a highly confident core prediction set.

### Implementing the LiPLike approach

Although LiPLike is available as a standalone GRNI tool, its underlying algorithm is remarkably straightforward. Given the expression matrices of transcription factors and target genes, LiPLike can be implemented in just six lines of Python code (see Code Listing 1), offering accessibility even to those with limited programming experience. Code Listing 1 demonstrates a direct implementation of Eq. 5 using Python 3.12 and NumPy 1.26, illustrating the core LiPLike procedure.

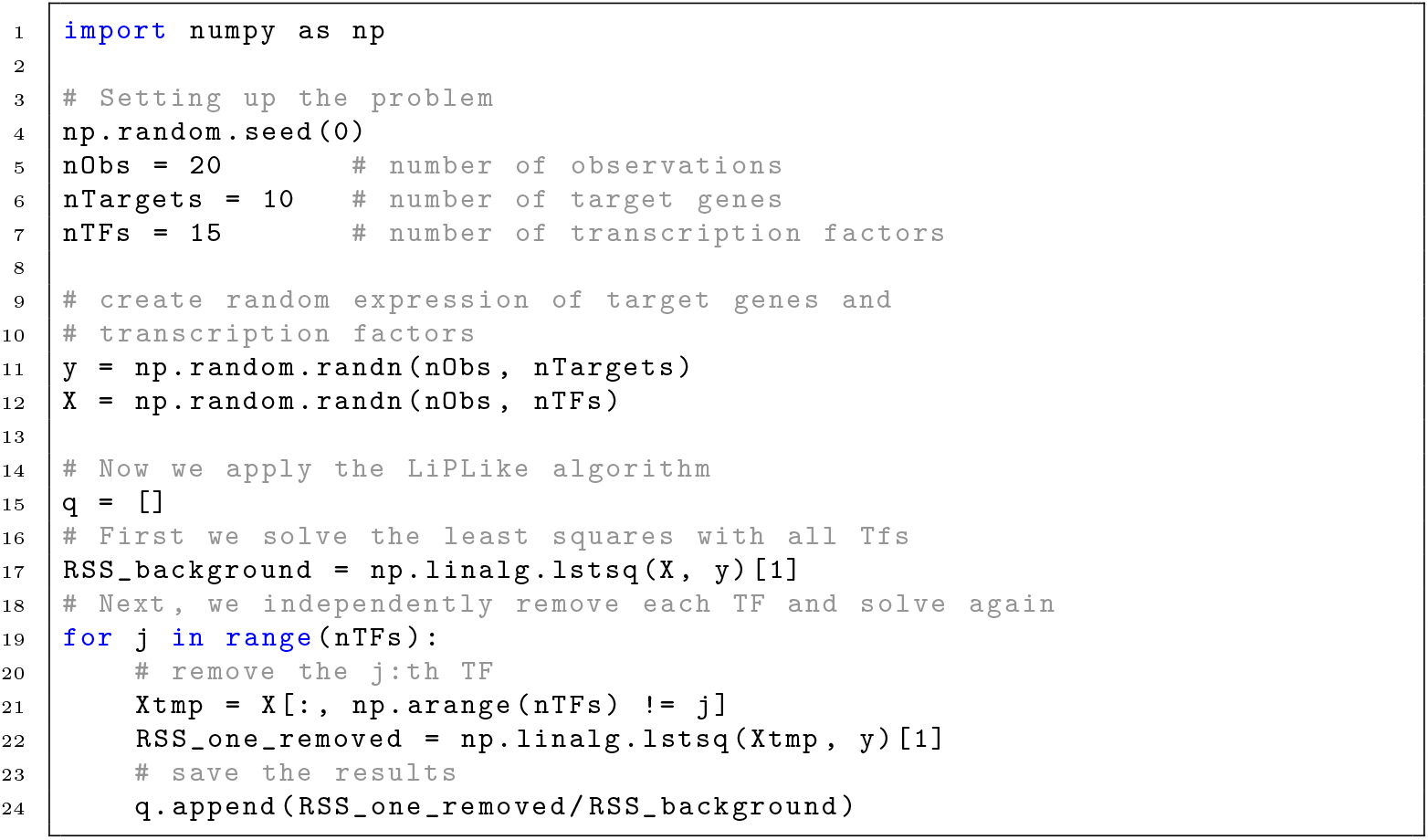

While Code Listing 1 is enough to implement LiPLike, a toolbox can be accessed at https://gitlab.com/Gustafsson-lab/liplike.

## 4 Notes

### Reworking LipLike for estimations of *q*_Δ_ and *q*

LiPLike ranks the confidence of predicted gene regulatory interactions using the statistic *q*, which compares the fit of a linear model with and without the specific interaction. Importantly, when the denominator in Eq. 6 approaches zero, even small changes in the numerator can disproportionately inflate *q*, potentially leading to overconfident predictions. To mitigate this issue, we introduce the statistic *q*_Δ_, as written in Eq. 8.

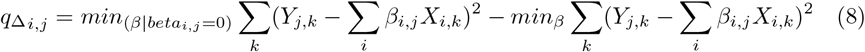

Equation 8 uses the same notation as in Eq. 6-7 but measures the difference between the two residual sum of squares (RSS) values, rather than their ratio. However, relying solely on this difference introduces a reverse issue: genes with large RSS values become disproportionately favored, potentially biasing the selection toward poorly fitting targets. o balance these effects, we recommend using a combined threshold based on both *q* and *q*_Δ_ (Fig. 6).

**Figure 6:**
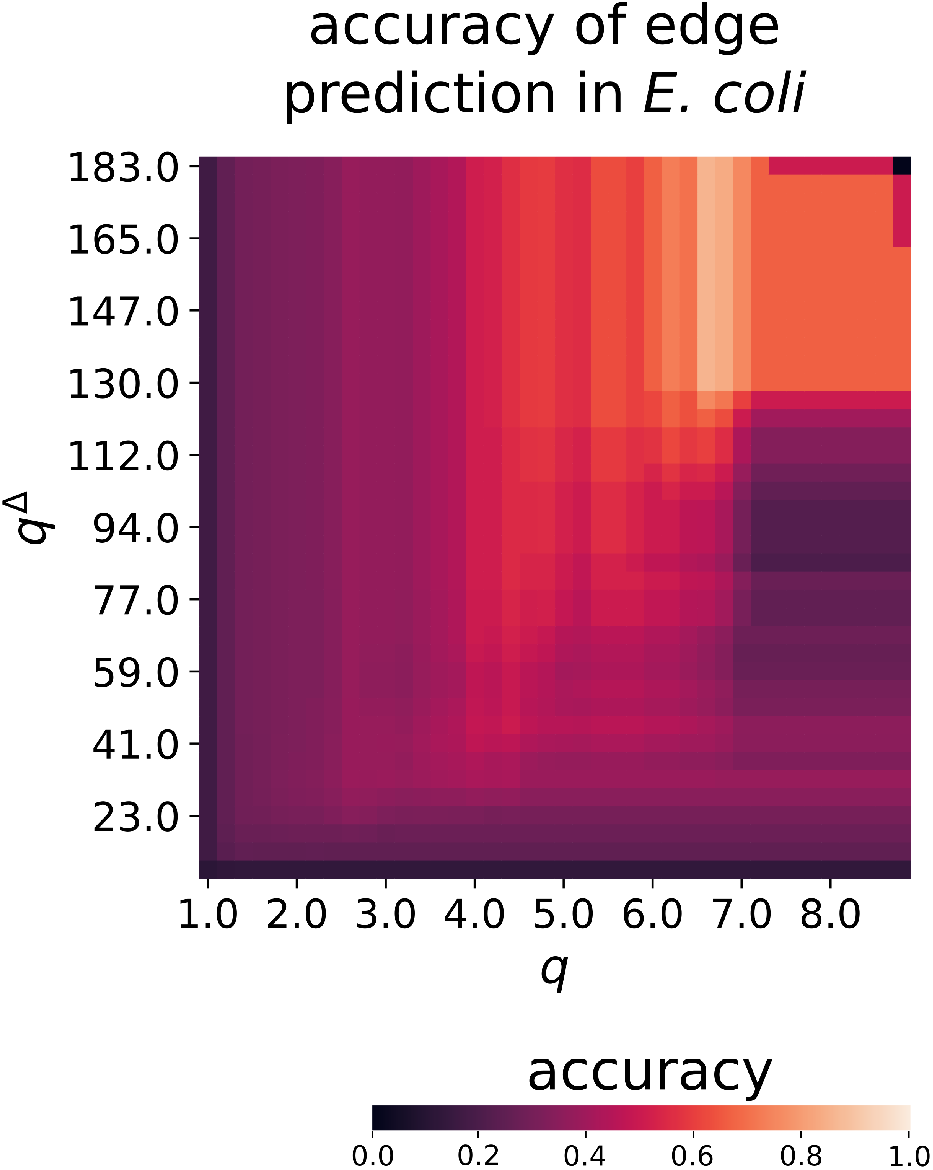
Prediction accuracy as a function of *q* and *q*_Δ_. The plot shows gene-gene regulatory prediction accuracy based on cutoffs of *q* and *q*_Δ_, using the Escherichia coli gene expression dataset from Marbach et al. (2012). While both metrics can independently reduce false positives, their combined use significantly improves performance. Figure adapted from Magnusson (2020) [14].

### Showcasing LiPLike applied to RNA-seq data from pancreatic cancer samples

To showcase how LiPLike behaves in GRNI, it was applied to gene expression data from primary tumors of pancreatic adenocarcinoma (PAAD), as made available by The Cancer Genome Atlas (TCGA). This dataset contains 183 samples, which was divided further into 23,360 target genes and 1,493 TFs, using the annotations presented in The Human Transcription Factors [10]. As the problem is underdetermined, LiPLike was implemented with an *L*_1_-penalty, with the regularization term 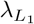 selected using cross-validation as per a standard LASSO setup.

LiPLike is fully parallelizable, with computational time in practice scaling linearly with the number of independent variables, i.e. TFs. The regularization type will set parameter values to exactly zero [25] during the estimation of the denominator of Eq. 6, which greatly increases computational speed, as most parameters will be estimated to a value of exactly 0 in the full network, leaving further analysis redundant. In this particular case, the LiPLike inference on average took 53 times longer per target gene than the LASSO equivalent.

Analysis of the top regulators in the LiPLike-inferred gene regulatory network (Fig. 7) identified *ZNF69* as the most prominent TF, with the twelve highest *q*-values all associated with it. The gene *ZNF69* regulates the transcription of RNA polymerase II, making it a highly interesting TF from a cancer perspective [24]. Other examples of TFs corresponding to the top *q*-values were *MEIS3*, which promotes the proliferation of pancreatic beta-cells [11], *ZNF559*; another RNA polymerase II regulator [20], and *NFE4*, which is believed to play a role in RNA polymerase II recruitment [29]. Importantly, the TF-target links identified as the most certain by LiPLike do not necessarily correspond to the most biologically impactful regulators. Rather, these high-confidence predictions should be interpreted as a core set of interactions, useful for refining predictions from alternative GRNI methods.

**Figure 7:**
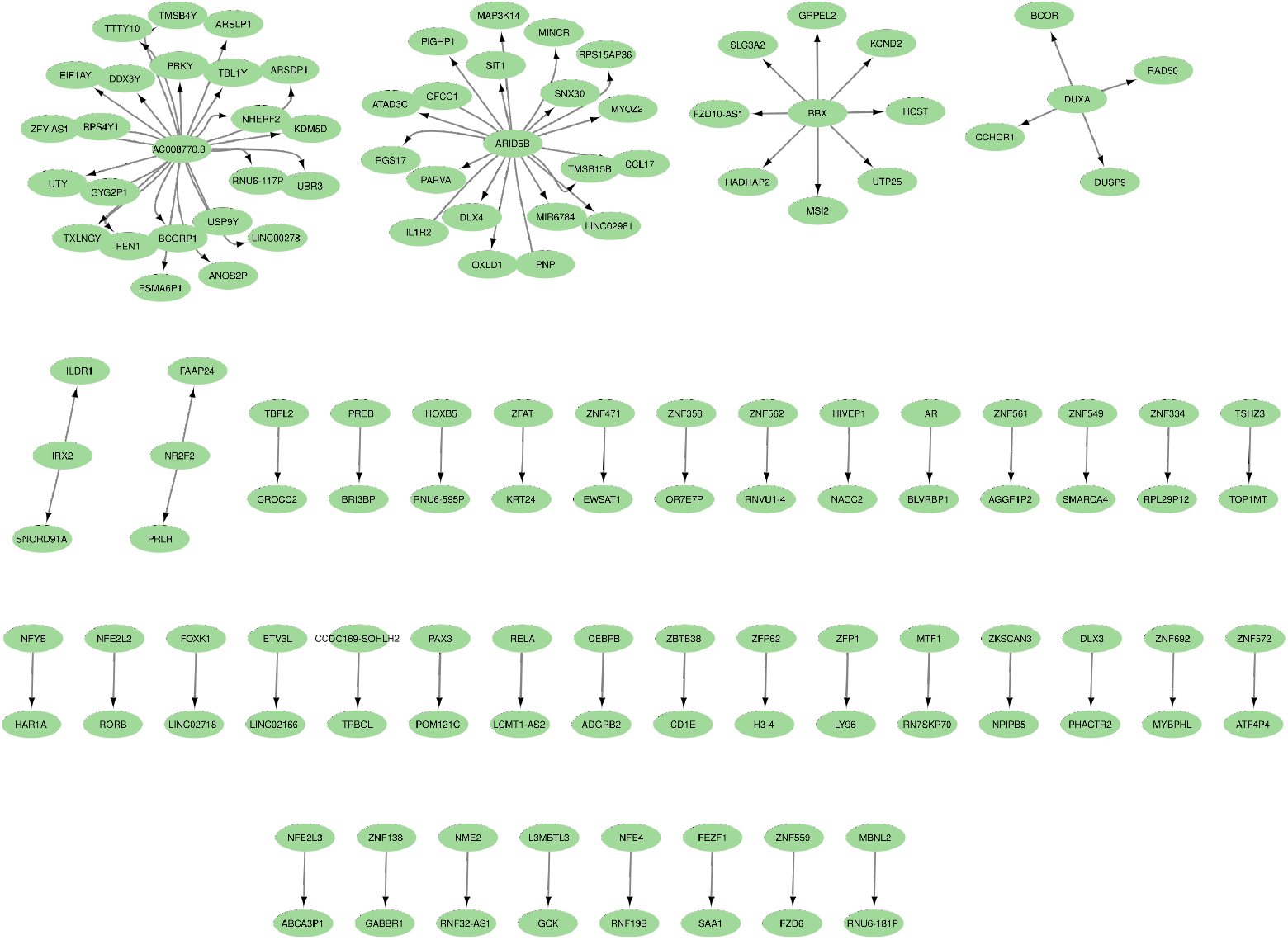
Network of irreplaceable TF-target interactions in PAAD data. Notably, the inferred network is considerably smaller than that of a typical GRNI tool, and should instead be viewed as a core or high-accuracy predictions.

A comparison between the LiPLike *q*-statistic and LASSO-derived coefficients revealed a significant correlation (Spearman *ρ* = 0.42, p *<* software detection limit; Fig. 8a). LiPLike analysis was only performed where the LASSO-estimated coefficient was non-zero. Most of the TF-target regulation predictions exhibited a low LiPLike *q*-value, suggesting them to be not uniquely determined from the data. Importantly, a limited subset of the regulation predictions was associated with substantially higher LiPLike *q*-values, suggesting them to be high confidence predictions (Fig. 8a). From a LASSO-centric perspective, the top predictions, here measured by the amplitude of the absolute value of the inferred *β* coefficients, were of mixed confidence as inferred by LiPLike. Consider the top TF-target regulation predictions, defined as where the absolute value of the inferred LASSO coefficient, |*β*_*LASSO*_|, was higher than ~0.75, as shown in Figure 8a. Approximately half of these interactions were about as linearly replaceable as the typical TF-target pair, as indicated by LiPLike, and are marked with a red background in Figure 8a. The reminder, however, had a disproportionately high *q*-value, as inferred by LiPLike, and should thus be considered top-confidence predictions.

**Figure 8:**
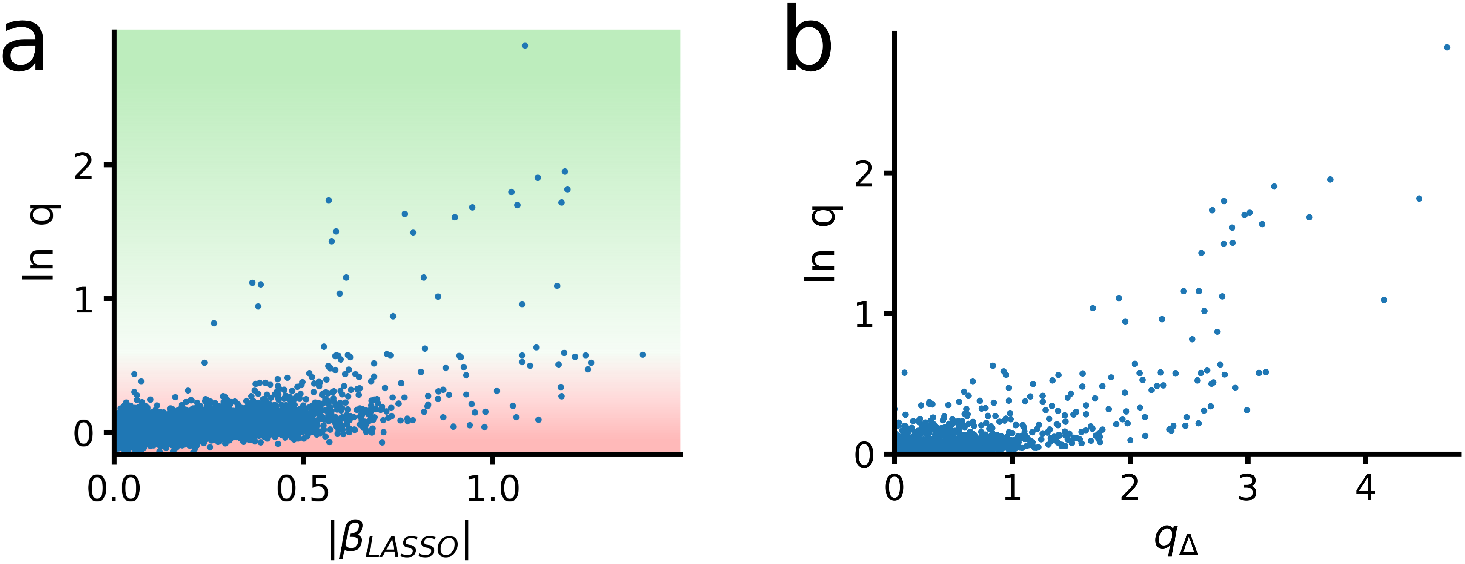
Case study of LiPLike applied to PAAD data. (a) LiPLike *q*-values (y-axis) vs. absolute LASSO-inferred TF-target coefficients (x-axis). Red background highlights interactions that are linearly replaceable; green indicates uniquely required interactions with high q-values. (b) Comparison of LiPLike metrics *q* (y-axis) and *q*_Δ_ (x-axis). While correlated, the metrics capture different aspects of irreplacability, with *q* offering finer discrimination among high-confidence predictions.

Further refinement of prediction accuracy was achieved using the alternative metric *q*_Δ_ (Eq. 7), defined as the residual difference between the full model and a model with one TF removed (Fig. 8b). This measure, called *q*_Δ_ in Eq. 8, was calculated along with the *q* metric when LiPLike was applied to the PAAD gene expression data (Fig. 8b). These two metrics were strongly correlated (Spearman *ρ* = 0.80, p *<* software detection limit), yet a subset of predictions with *q*_Δ_ *<* 1.5 had unexpectedly low *q*-values, suggesting *q* may provide a more robust separation of high-confidence predictions.

## 5 Concluding remarks

This chapter has presented the LiPLike approach for GRNI. The rationale behind LiPLike is to predict interactions with high probability, as opposed to recreating full networks. Importantly, LiPLike relies on either penalized or non-penalized linear regression models to estimate these uniquely needed interactions. However, biological systems are inherently non-linear, meaning LiPLike will never fully account for the possibility of alternative solutions in GRNI.

One of the main takeaways from the DREAM5 GRNI challenge was that an aggregate of inferred gene regulatory networks outperformed any individual GRNI method. It is therefore reasonable to speculate that the rationale behind LiPLike could extend beyond the linear domain. A logical continuation would involve using a compendium of GRNI methods, independently removing each regulator from the system, and comparing the results. Although such an approach would address some limitations of working in linear space, it would still be subject to biological constraints, such as using mRNA as a proxy for TF protein activity, despite the often low correlation between mRNA expression and explanatory power [1].

Nevertheless, the state-of-the-art in GRNI remains confined to a limited set of methods, most of which focus on predicting the most likely, complete gene regulatory network [22]. LiPLike can be used to stratify the gene-gene predictions of such networks into two sets: one comprising replaceable predictions and another containing linearly uniquely needed interactions with high confidence.

## Acknowledgments

This work was supported by the Swedish Research Council (grant no.: Dnr 2019-03767) and Stiftelsen Hierta Retzius stipendiefond (vetenskapliga ändamål), as administered by the Royal Swedish Academy of Sciences (BS2024-0053). The author thanks Associate Professor Elin Nyman for proofreading the manuscript.

